# IntroSpect: motif-guided immunopeptidome database building tool to improve the sensitivity of HLA binding peptide identification

**DOI:** 10.1101/2021.08.02.454768

**Authors:** Le Zhang, Geng Liu, Guixue Hou, Haitao Xiang, Xi Zhang, Ying Huang, Xiuqing Zhang, Bo Li, Leo J Lee

## Abstract

Although database search tools originally developed for shotgun proteome have been widely used in immunopeptidomic mass spectrometry identifications, they have been reported to achieve undesirably low sensitivities and/or high false positive rates as a result of the hugely inflated search space caused by the lack of specific enzymic digestions in immunopeptidome. To overcome such a problem, we have developed a motif-guided immunopeptidome database building tool named IntroSpect, which is designed to first learn the peptide motifs from high confidence hits in the initial search and then build a targeted database for refined search. Evaluated on three representative HLA class I datasets, IntroSpect can improve the sensitivity by an average of 80% comparing to conventional searches with unspecific digestions while maintaining a very high accuracy (∼96%) as confirmed by synthetic validation experiments. A distinct advantage of IntroSpect is that it does not depend on any external HLA data so that it performs equally well on both well-studied and poorly-studied HLA types, unlike a previously developed method SpectMHC. We have also designed IntroSpect to keep a global FDR that can be conveniently controlled, similar to conventional database search engines. Finally, we demonstrate the practical value of IntroSpect by discovering neoantigens from MS data directly. IntroSpect is freely available at https://github.com/BGI2016/IntroSpect.

## INTRODUCTION

The study of immunopeptidome, which is the collection of peptides presented on a cell surface by major histocompatibility complex (MHC) molecules, is invaluable to the development of next-generation vaccines and immunotherapies against autoimmunity, infectious diseases, and cancer [1-8]. Usually, the identification of immunopeptidome by mass spectrometry (MS) is carried out with standard database search tools [9,10], such as MS-GF+ [11], Comet [12], Open pFind [13], X!Tandem [14], and MaxQuant [15]. These tools, mainly designed for shotgun proteome, are not able to effectively handle the huge search space caused by unspecific digestion in immunopeptidome [16-18], resulting in low sensitivity and/or high false positive rates on peptide identification [19-24].

Conventional database search for immunopeptidome consists of the following steps: generating search space by unspecific digestion, assigning the spectra of MHC-bound peptides to their sequences, and scoring and filtering assignments by a certain false discovery rate (FDR) [25,26]. Unspecific digestion could potentially cut proteins between every two adjacent amino acids, leading to a huge search space. Previous studies have suggested that the overly inflated search space could over-estimate FDR and lead to low sensitivity [27-29]. To increase the sensitivity of immunopeptidome database search, two classes of computational methods have been developed: the first class, including MSrescue [22], DeepRescore [23], and MHCquant [30], aims to optimize the scoring and filtering of assignments and will be referred to as post-processing tools in this manuscript; the second class, such as SpectMHC [19], aims to optimize the generation of search space and will be referred to as database building tools. SpectMHC builds the targeted search space based on HLA-peptide binding predictions, which is trained from existing HLA-binding peptide databases. Its performance will be heavily influenced by the accuracy of the corresponding binding prediction, which may not work well for poorly-studied HLAs [22]. Furthermore, SpectMHC combines the iterative searches of unspecific digestion database and HLA-binding peptide database, making it infeasible to calculate a global FDR [31], which is important for controlling the overall error rate [32].

To increase the sensitivity of immunopeptidome detection, we developed a novel motif-guided immunopeptidome database building tool named IntroSpect. IntroSpect trains data-efficient PSSM models based on the high scoring peptides identified by conventional database search and builds a targeted database to carry out refined search. In the remainder of this paper, we will detail the development of IntroSpect, demonstrate its superior performance over existing database building tools and show how it can be used to identify neoantigens from MS data directly, an important application in cancer immunotherapies. We believe our freely available, open-source tool makes a significant contribution to advance the field of immunopeptidomics.

## MATERIALS AND METHODS

### Generation of cell lines

The K562 and HCT116 cell lines were obtained from ATCC (American Type Culture Collection, Manassas, VA), and the K562 cell line was engineered to express a single HLA-allele as follows. It was transduced using a highly efficient retroviral vector coding HLA-A*11:01. The vectors were transfected into a 293T packaging cell line and replication-defective virus supernatants were harvested. After infection of K562 cells with the supernatant, antibody-directed flow cytometry sorting was done to obtain cells with high expressions of HLA-A*11:01. Cells were grown in T75 flasks to a density of 1 × 10^9^ cells before harvesting for experiments.

### Purification of HLA-I peptides

HLA-I peptides were obtained from K562 and HCT116 cells as described previously [33]. In brief, 1 × 10^9^ cells were dissociated using 40 ml of lysis buffer with 0.25% Sodium deoxycholate, 1% n-octyl glucoside, 100 mM PMSF and protease inhibitor cocktails in PBS at 4 °C for 60 min. Lysate were further cleared by 30 min centrifugation at 14,000 g. Cleared lysate were immunoaffinity purified with pan-HLA class I complexes antibody covalently bound to Protein-A Sepharose CL-4B beads. Beads were first washed with 10 column volumes of 150 mM NaCl, 20 mM Tris HCl (buffer A), then 10 column volumes of 400 mM NaCl, 20 mM Tris HCl, then 10 volumes of buffer A again, and finally with 10 column volumes of 20 mM Tris HCl, pH 8.0. The HLA-I molecules were eluted at room temperature using 0.1 N acetic acid. Eluate were then loaded on Sep-Pak tC18 cartridges (Waters, 50mg) and washed with 0.1% TFA. The peptides were separated from HLA-I complexes on the C18 cartridges by eluting with 30% ACN in 0.1% TFA and concentrated to 20 µl using vacuum centrifugation. Finally, 5 µl sample was used for MS analysis.

### LC-MS/MS analysis of HLA-I peptides

HLA-I peptides of K562 and HCT116 cells were separated by HPLC (15 cm long, 75 µm inner diameter columns with ReproSil-Pur C18-AQ 1.9 µm resin) and eluted into an Orbitrap Fusion Lumos mass spectrometer (Proxeon Biosystems, Thermo Fisher Scientific). Peptides were separated with a gradient of 2–30% buffer (80% ACN and 0.5% acetic acid) at a flow rate of 250 nL/min over 65 min. MS was performed using data-dependent acquisition (DDA) mode. MS1 scans were conducted at a resolution of 120,000 over a scan range of 350-1500 m/z with a target value of 3 × 10^6^. Based on MS1 scans, MS2 scans were conducted at a resolution of 60,000 at 100 m/z with a target value of 1 × 10^5^. Fragment ion was produced by higher energy collisional dissociation (HCD) at 28% collision energy with a precursor isolation window of 2 m/z.

### Sequencing and analysis

For HCT116 cell line, DNA extractions, libraries construction, and sequencing (pair-end 100bp) were conducted according to protocols of MGISEQ-2000 platform (BGI-Shenzhen, China). RNA-Seq data of the HCT116 cell line were downloaded from the NCBI (SRR4228899). Low-quality reads were removed with SOAP nuke v1.5.6 [34]. DNA-Seq data were processed by minimap2 v2.11 [35] for read alignment and GATK [36] for variant analysis. RNA-Seq data were processed by HISAT v2.1.0 [37] for read alignment, GATK v3.7.0 [36] for variant analysis and RSEM v1.3.0 [38] for transcript quantification. A total of 480,905 potential neoantigens (9-11 mer) were generated from the sequences with variants.

### Mass spectrometry database search

MS data of B721.221 [39] and Jurkat [22] cell lines were downloaded from public databases (B721.221, MSV000080527 in MassIVE; Jurkat, PXD011723 in PRIDE). The raw files of public and inhouse MS data were converted to mgf files using ProteoWizard msConvertGUI [40]. The MS-GF+ search tool (release 2018.07.17) was employed to search the databases against the converted MS data. The conventional database contains 161,521 Uniprot [41] human protein entries (2017.12.20) and 245 frequently observed contaminants such as human keratins, bovine serum proteins, and proteases. For HCT116 dataset, 480,905 potential neoantigens were added to the database. Parameters of MS-GF+ are: variable modifications, N-terminal acetylation (42.010565 Da) and methionine oxidation (15.994915 Da); enzyme, unspecific cleavage; precursor ion tolerance, 10 ppm; peptide length, 9-11; charge, 2-5. For IntroSpect search, the database contains the peptides that passed the filtering of PSSM models and the peptides identified by conventional search. Except that the enzyme was set to no cleavage, other parameters were the same as those of conventional search. Percolator [42] (version 3.02.0) post-processing tool was applied for the estimation at the peptide level of <1% FDR for both IntroSpect and conventional search. From the pout.tab output file generated by Percolator, assignments to the contaminants were eliminated.

### Gibbs clustering of HLA-I peptides

The peptides identified by conventional database search was clustered into various groups using GibbsCluster-2.0 Server [43], with the following parameters: number of clusters, 1-6; motif length, 9; max deletion length 2; max insertion length 0; number of seeds for initial conditions, 5; penalty factor for inter-cluster similarity, 0.8; weight on small clusters, 5; use trash cluster to remove outliers, enable; threshold for discarding to trash, 2; number of iterations per sequence per temperature step, 10. The peptides in the clusters with the highest KLD were retained for further analysis.

### PSSM model training and filtering

Based on the clusters, we built PSSM models as described previously [44] to learn the corresponding sequence motifs for peptides in different groups. Briefly, each element *P*_*ai*_ in the PSSM matrix is the likelihood of a specific amino acid *a* at a given position *i*. We calculated *P*_*ai*_ as follows

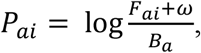

where *F*_*ai*_ denotes the frequency of a specific amino acid at the specific position in the peptides identified by conventional search; *B*_*a*_ denotes the frequency of the specific amino acid from a background database (such as Uniprot human protein database); and *ω* is a random value (ranging from 0 to 1) generated from a Dirichlet distribution [45]. To filter the whole proteome database to generate a targeted one, we define the motif score of a given peptide as the sum of the *P*_*ai*_ at each site in the PSSM and only kept those with a motif score greater than 0.3.

### Synthetic peptide validation

A total of 118 randomly selected peptides from K562 dataset were synthesized and analyzed under the same MS conditions with K562 HLA I peptides. The mirror plots of spectra between synthetic peptides and eluted peptides were generated by PDV [46]. To validate a peptide which could be presented by MHC-I complex, the following criteria were considered: i) the variation of retention time between precursor ions was less than 3 min; ii) the pattern and retention time were matched between synthetic and target peptides with no less than 5 product ions.

### Peptide Pearson Correlation Coefficient (PCC) calculation

To quantify the similarity between two sets of peptides with the same length, we calculated the Pearson Correlation Coefficient (PCC) of the amino acid frequencies between them. For a given position *i*, we first calculated the empirical probability mass functions (pmfs) of the amino acid distributions in both the first (*x*) and second (*y*) sets. The PCC between these two random variables *X*_*i*_ and *Y*_*i*_, *PCC*_*XiYi*_, is then computed as

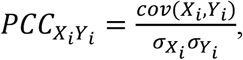

where cov is the covariance and *σ’*s are the standard deviations.

### Code Availability

We have made IntroSpect available on GitHub: https://github.com/BGI2016/IntroSpect. It is a command-line tool written in Perl which requires GibbsCluster v2.0 preinstalled, in Darwin (Mac) or Linux platforms. The tool takes an input protein FASTA database and peptides identified by conventional search and outputs targeted database which could be used for refined high sensitivity identification.

## RESULTS

### The development of IntroSpect

In order to reduce the overly inflated search space caused by unspecific digestions, we developed a strategy of motif-guided digestion in IntroSpect. The motif-guided digestion leads to a small and targeted database in which the peptides that are extremely unlikely to be present in a given sample will be filtered out. Peptides that do exist in the sample will get higher q values due to less competitions, making it easier for real peptides to stay after FDR filtering. Therefore, IntroSpect can achieve higher statistical power, and identify more peptides at the same FDR.

Search with IntroSpect includes four steps (Figure 1a). Step 1 is to import the conventional protein database and MS raw data into the search engine, and obtain peptides that pass 1% FDR filtering. These high-confidence peptides are then clustered into groups by GibbsCluster2.0 in step 2, and peptides in the same group are used to train a position-specific scoring matrix (PSSM) model to learn their motifs. In step 3, the PSSM model is used to score each peptide in the conventional database, and peptides with PSSM score > 0.3 (the optimal threshold in our test) as well as those with FDR < 1% in the first-round are combined to become the new search space. Step 4 runs the second-round search against this new, targeted database to identify peptides that pass 1% FDR as the final output. Unlike previous multi-round search strategies [19-21] where different rounds of results are combined, we decide to add the first-round peptides directly into the targeted database for the second (and final) round search, so that a global FDR can be obtained. As we will show later, the vast majority of first round peptides will still appear in the final results. Steps 2 and 3 are written in Perl to form the IntroSpect package, while steps 1 and 4 are left to the users to decide how to build the conventional database and to run the search engine of their choice.

**Figure 1.**
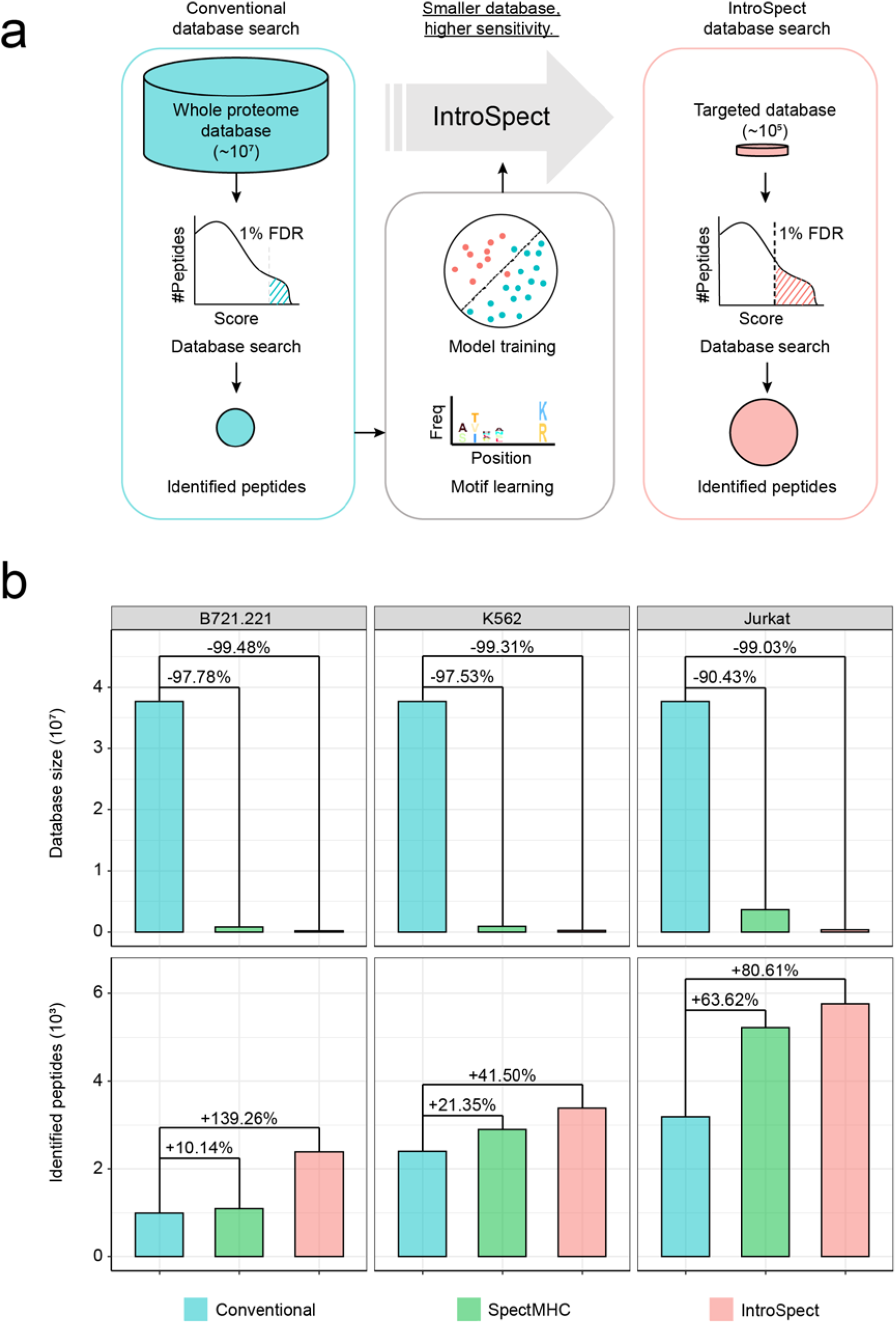
IntroSpect improves peptide identification sensitivity by reducing the search space. (a) The flowchart of the conventional database search and IntroSpect database search. (b) IntroSpect significantly decreases the database size and increases the identified peptides. The database size is calculated as the number of 9-11mer peptides in the database. The gray boxes on the top denote the cell lines.

### IntroSpect can identify substantially more peptides

To evaluate IntroSpect, we tested its performance on MHC class I immunopeptidome datasets from Jurkat, K562 and B721.221 cell lines (Table 1). We first used MS-GF+ as the search engine and Percolator for 1% FDR filtering, and ran IntroSpect, SpectMHC and conventional database search to identify 9-11mer peptides on these datasets [47,48]. The databases generated by Introspect are much smaller than the conventional databases (0.5%, 0.7% and 1.0% respectively, Figure 1b), and search with these reduced databases resulted in substantially more peptides under the same FDR (1.8-, 1.4- and 2.4-folds respectively, Figure 1b, Figure S1). The databases generated by Introspect are also significantly smaller than those by SpectMHC (∼1/4 to 1/10, Figure 1b), and the newly identified peptides are 1.3, 1.9, and 13.8 folds than those of SpectMHC (Figure 1b). We further calculated the ratio of assigned spectra to total spectra in these three search strategies. IntroSpect can assign 41.67%∼156.18% more spectra than those of conventional database search and 11.53%∼79.21% more than SpectMHC (Figure S2).

**Table 1.**
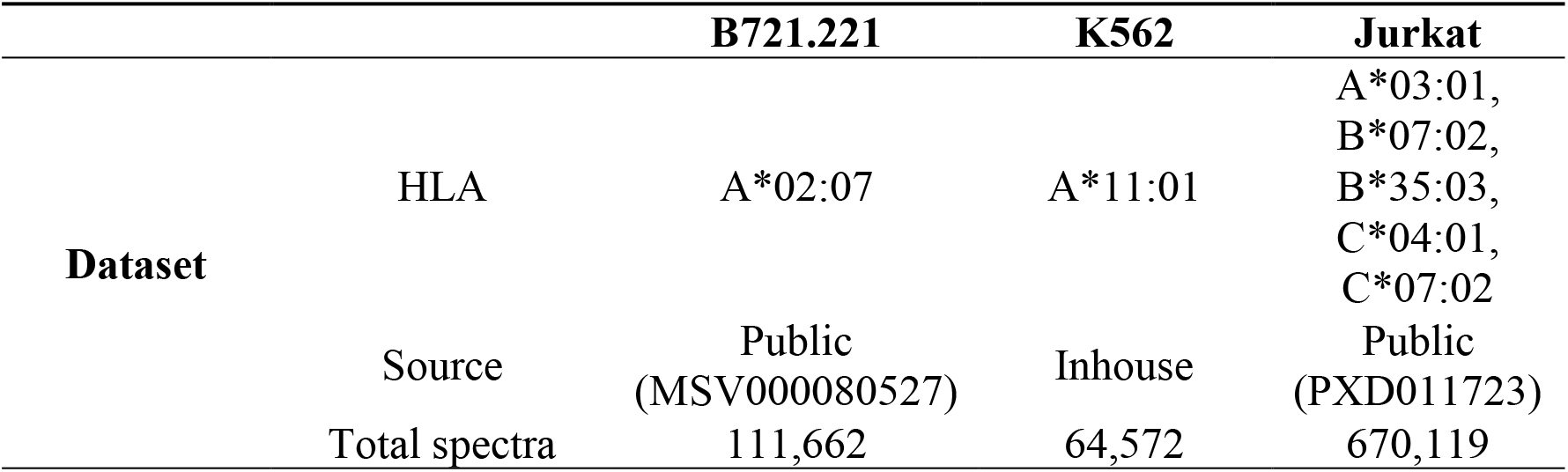
Summary of immunopeptidome data sets.

|  |  | **B721.221** | **K562** | **Jurkat** |
| --- | --- | --- | --- | --- |
| **Dataset** | HLA | A*02:07 | A*11:01 | A*03:01,<br>B*07:02,<br>B*35:03,<br>C*04:01,<br>C*07:02 |
|  | Source | Public<br>(MSV000080527) | Inhouse | Public<br>(PXD011723) |
|  | Total spectra | 111,662 | 64,572 | 670,119 |

In addition, we tested IntroSpect with another two common search engines, i.e., Comet (combined with Percolator) and MaxQuant, and observed the same trend. More specifically, when using Comet, IntroSpect can identify 3.3, 3.5 and 5.8 folds of total peptides comparing with conventional search and 1.0, 2.3 and 2.0 folds of new peptides comparing with SpectMHC; when using MaxQuant, the improvements are 2.9, 1.8 and 1.8 folds for conventional search and 2.6, 1.5 and 1.8 folds for SpectMHC respectively (Figure S3). These results clearly show that IntroSpect is not only much more sensitive than conventional search but also considerably more sensitive than SpectMHC, a previously developed method that also aims to optimize the search space for immunopeptidome.

### IntroSpect achieved a similar accuracy as conventional search

To assess the accuracy of peptides identified by IntroSpect, we first compared the proportion of identified peptides predicted to be binders by both IntroSpect and the conventional search. Similar strategies have been previously applied to check for the quality of MS data [22, 23, 39, 49]. We predicted IC50 values of peptides using netMHCpan 4.0 [50] and drew the distribution of IC 50 values for all identified peptides, with a vertical line indicating the cutoff for binders (IC50 < 500nm, Figure 2a). Please note that SpectMHC was not included in this analysis since netMHCpan has already been used when building the targeted database. We found that overall these two sets of distributions are very similar across the three datasets, with those identified by IntroSpect had slightly more binders (87.86% vs. 85.09% for Jurkat, 89.04% vs. 88.80% for K562 and 26.02% vs. 25.50% for B721.221). Notably, the percentage of predicted binders in the engineered single-allele B721.221 cell line, whose HLA type is A*02:07, is very low, as has been previously shown [39]. This is because A*02:07 is a poorly studied HLA allele, with a total of 180 binding affinity data points available, among which 22 are binders. This also partially explains why IntroSpec’s improvement over SpectMHC is particularly high for this dataset.

**Figure 2.**
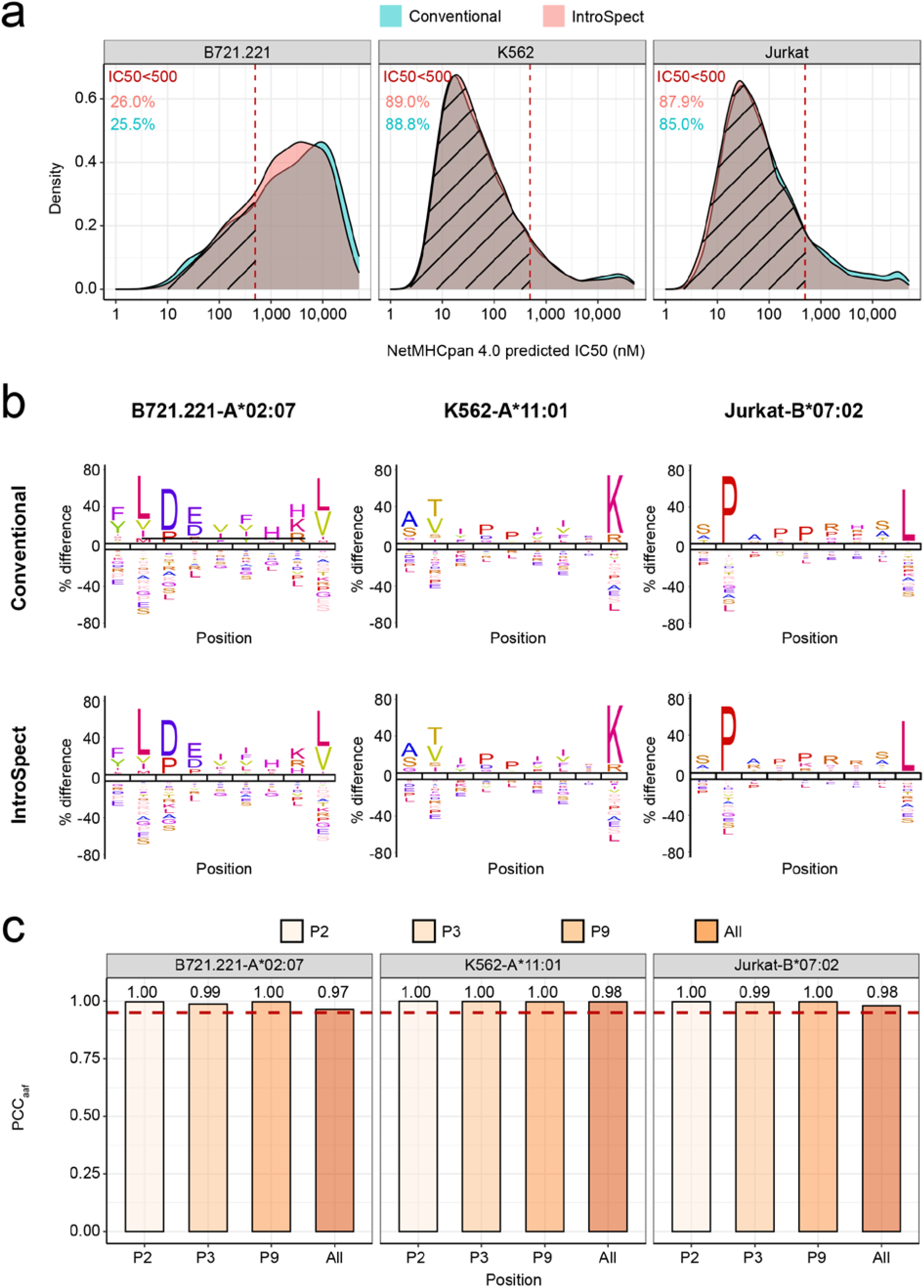
Immunopeptides from IntroSpect and conventional database search are very similar. (a) The distribution of predicted IC50 values of peptides identified by the conventional and IntroSpect search: the red dotted lines represent IC50 = 500nM, and the peptides in shaded area are predicted to be strong or weak binders, with their percentages marked above the figure. (b) The sequence logo comparison of immunopeptides in various datasets (B721.221-A*02:07, K562-A*11:01 and Jurkat-B*07:02) by the conventional and IntroSpect search. (c) Pearson correlation coefficients between the amino acid frequencies by the conventional and IntroSpect search at P2, P3, P9 and all positions of 9-mer peptides, the red broken line indicates 0.95.

We further visualized HLA-binding motifs using iceLogo [51,52]. Sequence logos of the representative 9-mer peptides from IntroSpect and conventional database search displayed high similarity in all datasets (Figure 2b). In addition, we quantified the similarities of the HLA-binding motifs based on the Pearson Correlation Coefficients between the amino acid frequencies (PCC_aaf_) of peptides of all positions. The average PCC_aaf_ of all positions (All) and each anchor position (P2, P3, P9) [53-56] are all greater than 0.95 (Figure 2c). We also obtained peptides of the corresponding HLA allele from IEDB and compared them with those obtained by us, and the results showed that the sequence motifs of our datasets were highly consistent with those from IEDB (Figure S4).

Finally, to validate the peptides identified by IntroSpect, we randomly selected a list of peptides from the K562 dataset. A total of 118 peptides (27 peptides newly identified by IntroSpect, 91 peptides identified by both methods) were synthesized and analyzed under the same MS acquisition conditions as that of K562 cell line. The spectra of synthetic peptides with highest PSM scores were then compared to the spectra of eluted peptides from K562 cell line in the experiment to confirm or reject the peptide identity. We found that 97.80% of the peptides (89 out of 91) identified by both methods could be confirmed by spectral validation, as well as 96.30% (26 out of 27) detected by IntroSpect method only (Table 2). Collectively, these results demonstrated that IntroSpect can not only identify many more peptides but also achieve an accuracy that is on a par with the conventional search method.

**Table 2.**
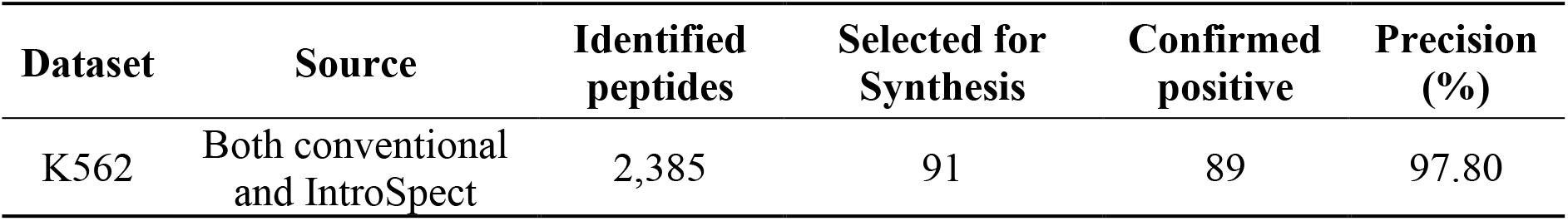
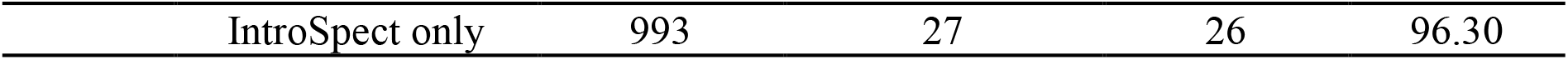
Randomly selected peptides identified by IntroSpect and conventional database search were confirmed by spectral validation

### IntroSpect inherits the results of conventinal database search

In MS data analysis, spectra provide the raw evidence for identified peptides. Therefore, the essence of newly identified peptides by IntroSpect is a reassignment of the spectra not recognized in conventional search. Based on IntroSpect’s methodology, we hypothesized that the identified spectra and peptides from IntroSpect would cover the vast majority of those from conventional search. Indeed, when we calculated the overlap of both identified spectra and peptides from the two methods, the overlapped spectra or peptides accounted for more than 99% of those identified by the conventional method in all three datasets (Figure 3a). Moreover, there were on average 48% of spectra and 44% of peptides identified by IntroSpect alone. We further observed that part of the unique spectra identified by IntroSpect (6% to 58%) matched to peptides already identified by conventional search, boosting the evidence of these previously identified peptides (17% to 88%, Figure 3b). We call them *refined peptides*, which are those that can be identified in the conventional search but assigned extra spectra by IntroSpect. It is worth pointing out that on average 21% of refined peptides had only 1 supported spectrum in the conventional search, but these proportions went down to zero with the added spectra identified by IntroSpect, while the proportion of refined peptides matched with more than 2 spectra increased from 58% to 90% (Figure 3c). Both lines of evidence, i.e., the overlap between IntroSpect and conventional search and the added support of IntroSpect identified spectra for refined peptides, showed the high consistency between these two search strategies, and validated our design choice of not simply aggregating different rounds of iterative search, which rendered the extra benefit of a unified global FDR.

**Figure 3.**
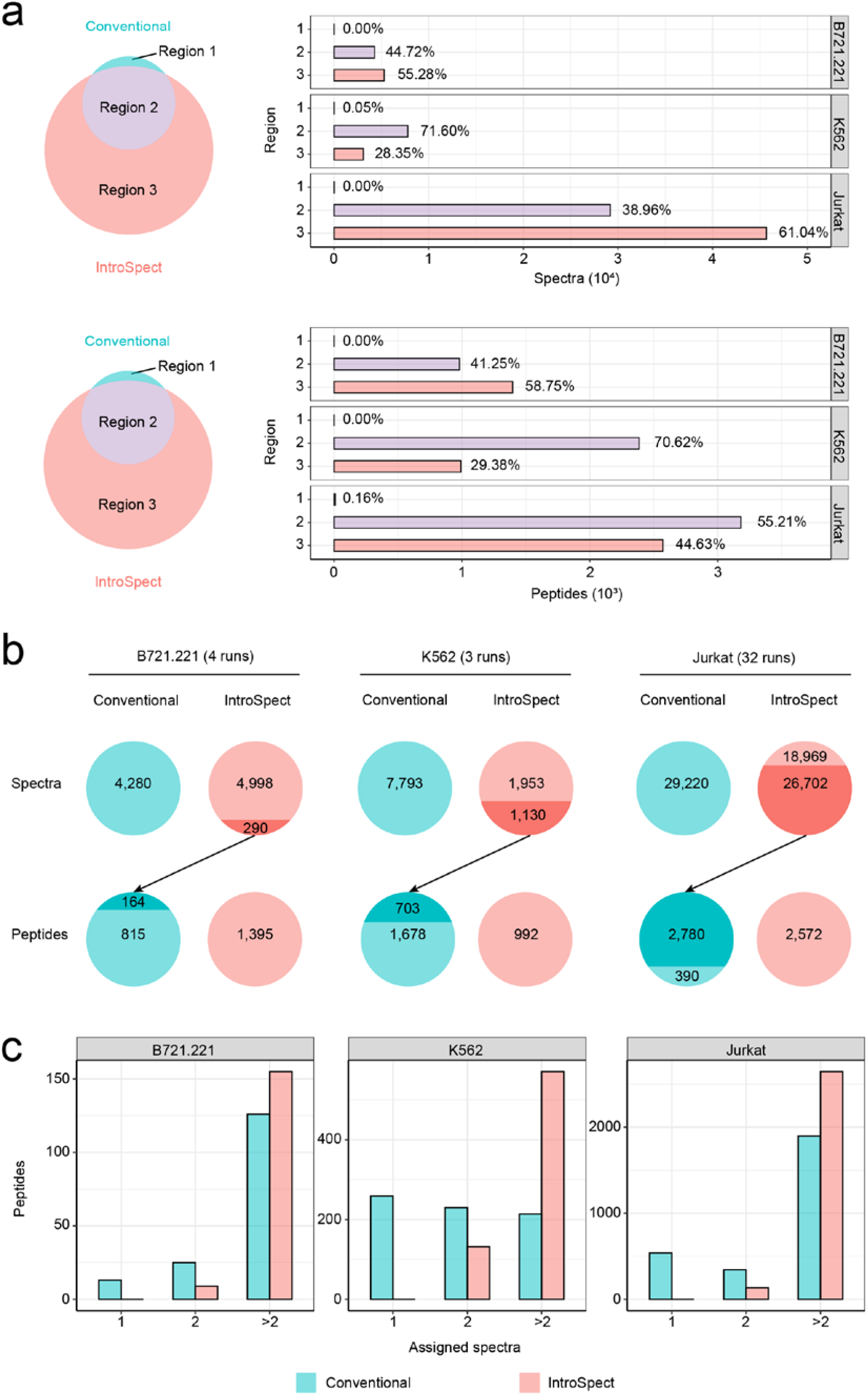
The high consistency in the spectra and peptides between the conventional search and IntroSpect search. (a) Most of spectra (top panel) or immunopeptides (bottom panel), detected by conventional method, can be identified through IntroSpect. Regions 1, 2, and 3 denote spectra (top panel) or immunopeptides (bottom panel), detected by conventional method only, both methods, or IntroSpect only. The percentages are calculated based on the total number of peptides or spectra identified by both methods. The gray boxes on the right panel denote cell lines. (b) A fraction of spectra newly identified by the IntroSpect matched to peptides (refined peptides) that had been previously identified by the conventional searching. The refined peptides were marked in dark color. (c) The assigned spectra for refined peptides were significantly increased.

### The database generated by IntroSpect is smaller and more targeted

Previous studies have suggested that small, targeted databases are beneficial for MS database search [16]. Here we have shown that IntroSpect does have a smaller database and is more sensitive than SpectMHC (Figure 1b, 1c). However, since IntroSpect learns motifs from the initial search results while SpectMHC learns motifs from external data, we suspect that their targeted databases differ more than just size. To investigate this, we adjusted the thresholds of IntroSpect and SpectMHC to obtain pairs of target databases with the same size for the K562 cell line, which has been engineered to express a single HLA-A*11:01 allele. All the generated databases of different sizes were used to identify peptides for the K562 dataset, and IntroSpect still had significant advantages over SpectMHC in terms of the numbers of identified peptides (Figure 4a). Furthermore, although the overlap between the databases by the two methods was small (∼20%), the overlap between the identified peptides was large (∼80%) and the numbers of peptides solely identified by IntroSpect was about 10 times more than those by SpectMHC across different database sizes (Figure 4b). Clearly, these results indicate that the database generated by IntroSpect is more targeted, or of higher quality when used in MS database search, comparing with that by SpectMHC. This is likely because motifs learned from the same MS data (as in IntroSpect) are a better match than those learned from external data (as in SpectMHC). To quantify, we calculated the average PCC_aaf_ at all positions between the peptides in the databases and those identified by SpectMHC or IntroSpect, and IntroSpect does have higher PCC_aaf_’s across different database sizes (Figure 4c, Figure S5). We also computed the same quantity across all three datasets with the default thresholds (PSSM score > 0.3 for IntroSpect and NetMHCpan 4.0 rank <2% for SpectMHC) of SpectMHC and IntroSpect, and observed the same trend (Figure 4d).

**Figure 4.**
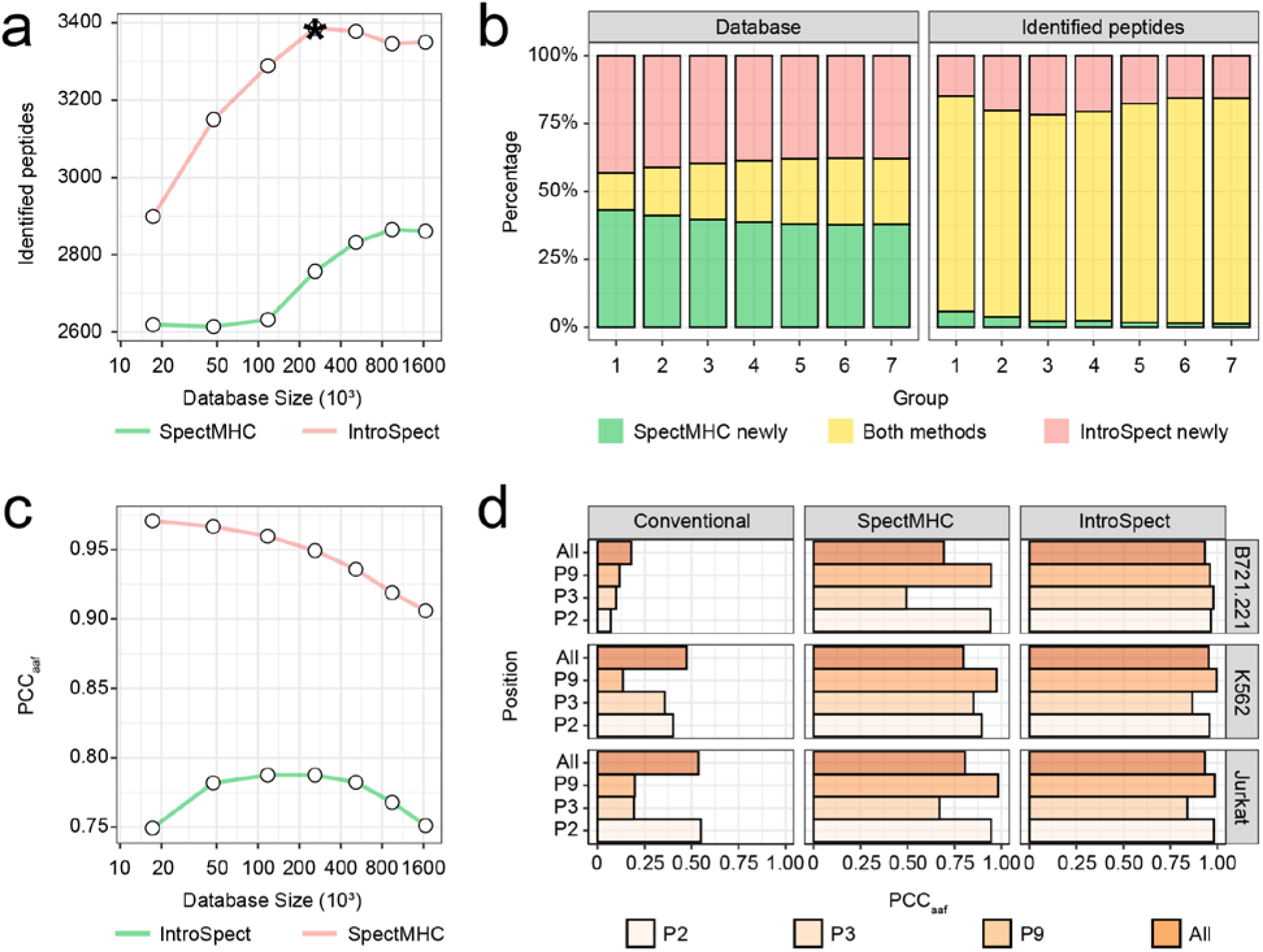
The IntroSpect generates a smaller and more targeted database than that of SpectMHC. (a) The line plot compared the numbers of identified peptides by SpectMHC and IntroSpect on databases with various matching sizes. The data point with an asterisk corresponds to the motif score of 0.3, the empirically chosen optimal threshold for IntroSpect. (b) The bar plot shows the relationship between the databases and identified peptides of IntroSpect and SpectMHC on databases with various sizes. (c) The line plot compared the PCC_aaf_ by SpectMHC and IntroSpect search on databases with various sizes. (d) PCC_aaf_ at P2, P3, P9 and P_aver_ positions of the database and identified peptides by SpectMHC, IntroSpect and conventional search on three datasets (B721.221, K562 and Jurkat).

### IntroSpect identified more neoantigens than conventional method

Having established the superior performance of IntroSpect, we next applied it to a key application in immunology, which is to directly identify neoantigens from MS profiling of the immunopeptidome. This is a very challenging problem since neoantigens are typically of low abundance. However, due to the practical importance of neoantigens in cancer immunotherapies, great efforts have been made to identify them in the past, going beyond the standard MS techniques, such as manual inspections of MS spectra without stringent FDR filtering [57] or experimentally altering the antigen processing machinery (APM) components to increase the abundance of neoantigens [58-61].

Here, we generated immunopeptidome as well as sequencing data for the HCT116 cell line by standard experimental techniques and focused on comparing the abilities of conventional search and IntroSpect in identifying neoantigens. Based on the sequencing data of HCT116, we first generated all 9-11 mer potential neoantigens and added them to the Uniprot database, and performed conventional and IntroSpect search as described previously. As before, IntroSpect was able to identify substantially more peptides than conventional search (2,222 versus 1,435), but more importantly, 10 neoantigens were identified by IntroSpect versus 4 by conventional search, a more than two fold increase (Figure 5a, Table S1). As expected, the q-values of these 10 neoantigens were significantly reduced in IntroSpect comparing with conventional search (Figure 5b). We also manually inspected the supported spectra of these 10 neoantigens and they are all of high quality (Figure 5c, Figure S6). To further exam the quality of these identified neoantigens, we exhaustedly search for established experimental evidence of them, including ligand presentation, qualitative binding, IFNg release assay etc. [62]. We were able to find previous evidence for 1 of the 4 neoantigens identified by conventional search but 5 of the 6 additional neoantigens discovered by InstroSpect (6 of 10 in total). Becker et al. recently proposed to use 5AZA to treat the HCT116 cell line to enhance its antigen presentation process and identified a number of extra neoantigens based on this technique [61]. Interestingly, while conventional search with our data was not able to identify any of the neoantigens discovered by Becker et al., IntroSpect was able to identify two of them (SLMEQIPHL and QTDQMVFNTY). In summary, we believe that this brief case study has shown the good potential of IntroSpect in practical applications.

**Figure 5.**
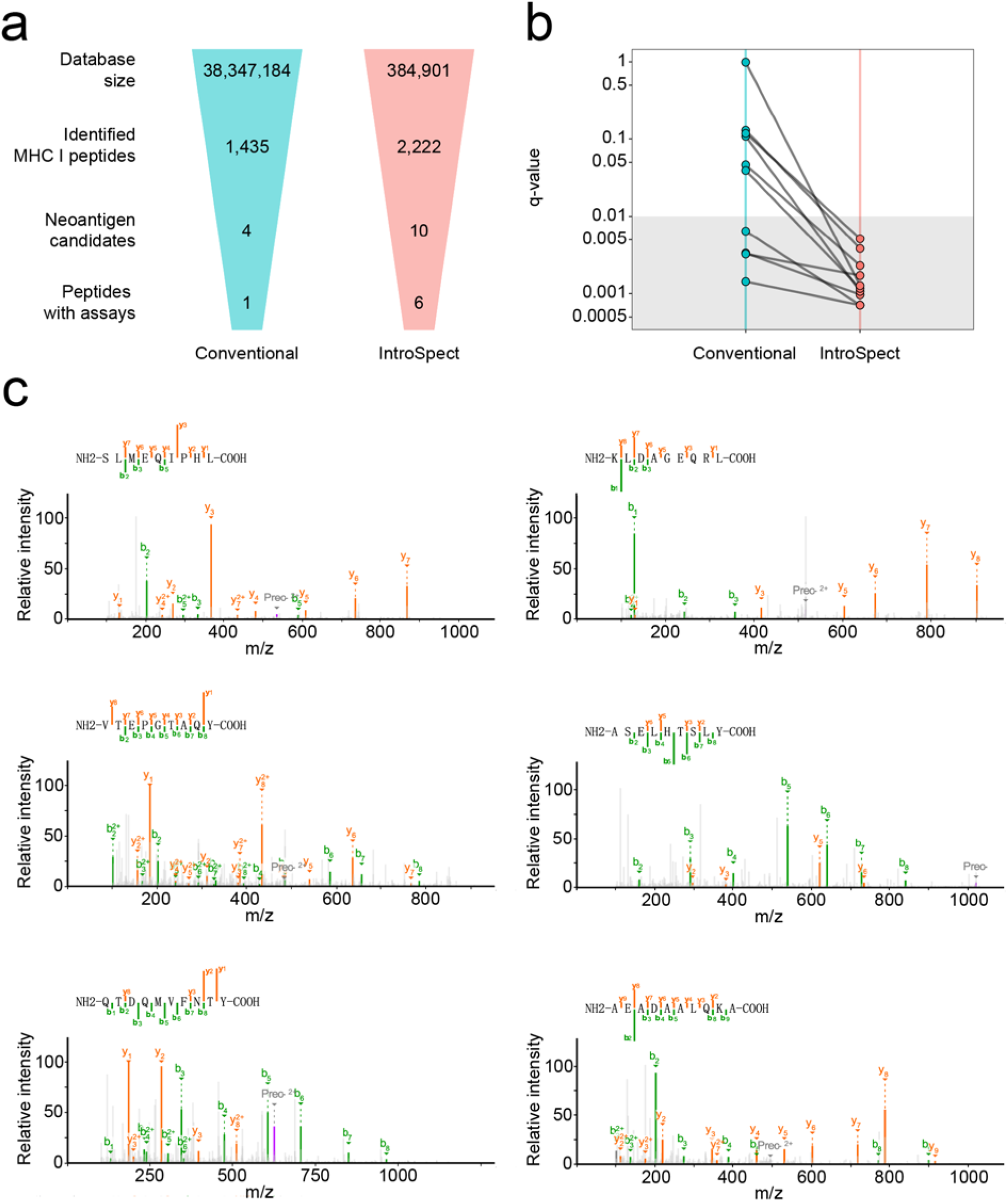
IntroSpect identified more neoantigens than the conventional search. (a) Flowcharts indicating key steps involved in neoantigen discovery. (b) Percolator q-values of neoantigens identified by both methods are plotted. The grids at the bottom of the plot mark the neoantigens with positive assay results from other research. The Neo-10 peptides (AEADAALQKA) has no q-value in the conventional search, and we define it to be 1. (c) Spectra of neoantigen candidates newly assigned by IntroSpect. Peaks represent b ions in green, y ions in orange and precursor ions in dark grey.

## DISCUSSION

Currently, high-throughput immunopeptidome profiling are usually based on MS database search, but the lack of specific digestion leads to low sensitivity. Here, we developed IntroSpect, a motif-guided immunopeptidome database building tool, to overcome this challenge. By testing on diverse immunopeptidome datasets, we showed that IntroSpect could significantly increase the sensitivity of identification comparing with not only conventional search, but also a previously developed database building tool SpectMHC, while maintaining a high accuracy. It is also worth mentioning that it can be easily combined with existing post processing tool as well to potentially achieve further performance improvement. However, IntroSpect is not without limitations. For example, the current PSSM model is peptide length and HLA allele specific, which means that the high-confidence peptides identified in the initial search must be further subdivided for model training. When the peptides identified from conventional search is relatively few, say <500, the training set of a certain length and HLA allele might be too small to effectively train the corresponding PSSM model, and in such cases, the external data based SpectMHC could perform better. We plan to next adopt deep learning techniques to leverage existing, large scale MS data to pre-train length independent sequence models and then adapt the pre-trained models to specific experiments by transfer learning. The motif scores, which only serve as an empirically chosen threshold to filter out highly unlikely peptides, could also be better utilized. One way to do so is to assign weighted prior probabilities for different peptides based on their motif scores when doing database search, similar to what has been done in Li S et al.’s recently developed constrained de novo sequencing approach [63]. Nonetheless, we believe the simple and effective strategy implemented in IntroSpect has significantly moved the quality of MS profiled immunopeptidome analysis forward and opened the door to apply this exciting MS technique in broader scenarios, such as in understanding non-canonical or post translationally modified immunopeptides [64,65].

## ASSOCIATED CONTENT

### Supporting Information

The following files are available free of charge. Supplementary Figure 1. The Q value distribution. Supplementary Figure 2. The percentage of spectra assignments. Supplementary Figure 3. The performance of IntroSpect on other search tools. Supplementary Figure 4. The PCC_aaf_ on each position. Supplementary Figure 5. The sequence logo comparison. Supplementary Figure 6. Spectra of neoantigens. Supplementary Table 1. The neoantigens identified from HCT116 cell line. (PDF)

### Data Availability

The MS data of K562 and HCT116 datasets have been deposited in the public proteomics repository MassIVE (https://massive.ucsd.edu) with accession number MSV000086567 and MSV000087927. The sequencing data as well as the above MS data have also been deposited into CNGB Sequence Archive (CNSA) [66] of China National GeneBank DataBase (CNGBdb) [67] with accession number CNP0001446.

## AUTHOR INFORMATION

### Author Contributions

Conceptualization, Leo J Lee and Le Zhang; methodology, Geng Liu, Guixue Hou and Ying Huang; software, Le Zhang and Haitao Xiang; validation, Le Zhang; formal analysis, Le Zhang; resources, Bo Li and Xi Zhang; writing, Le Zhang, Leo J Lee and Bo Li; supervision, Leo J Lee, Bo Li and Xiuqing Zhang.

### Notes

All authors are BGI employees.

## ACKNOWLEDGMENT

We thank the China National Gene Bank (CNGB) for its support in the sample sequencing and data storage.

## ABBREVIATIONS

HLA: human leukocyte antigen
MS: mass spectrometry
PSSM: position score specific matrixes
LR: logistic regression; major histocompatibility complex

