## Supplemental files for "IntroSpect: motif-guided immunopeptidome database building tool to improve the sensitivity of HLA binding peptide identification"

*Supplementary material lists*

Supplementary Figure 1. The Q value distribution of the conventional search and IntroSpect search.

Supplementary Figure 2. The percentage of spectra assignments.

Supplementary Figure 3. The performance of IntroSpect on Comet and MaxQuant search tools.

Supplementary Figure 4. The comparison of PCCaaf on each position between the IntroSpect and SpectMHC.

Supplementary Figure 5. The sequence logo comparison of immunopeptides in various datasets (B721.221-A*02:07, K562-A*11:01 and Jurkat-A*03:01) by the conventional search, IntroSpect search and IEDB downloaded.

Supplementary Figure 6. Spectra of neoantigen candidates assigned by both the conventional search and IntroSpect search.

Supplementary Table 1. The neoantigens identified from HCT116 cell line.

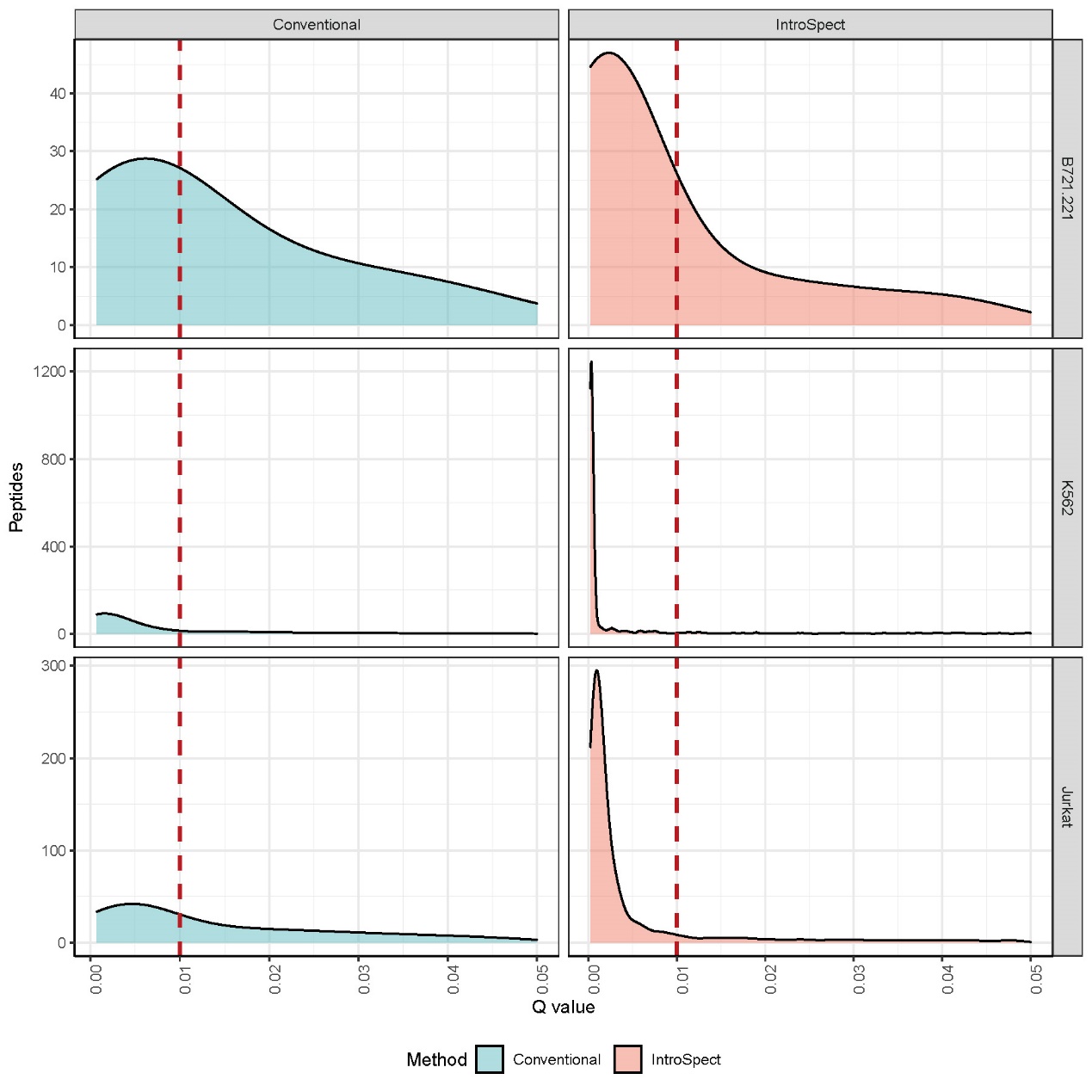

**Supplementary Figure 1.** The Q value distribution of the conventional search and IntroSpect search.

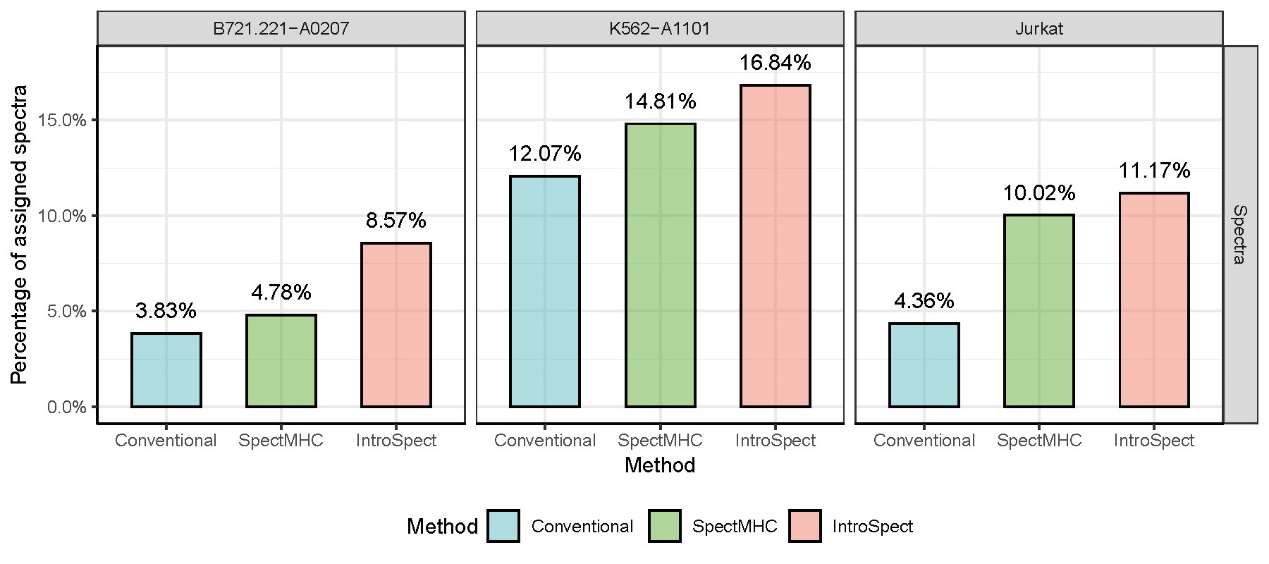

**Supplementary Figure 2.** The percentage of spectra assignments.

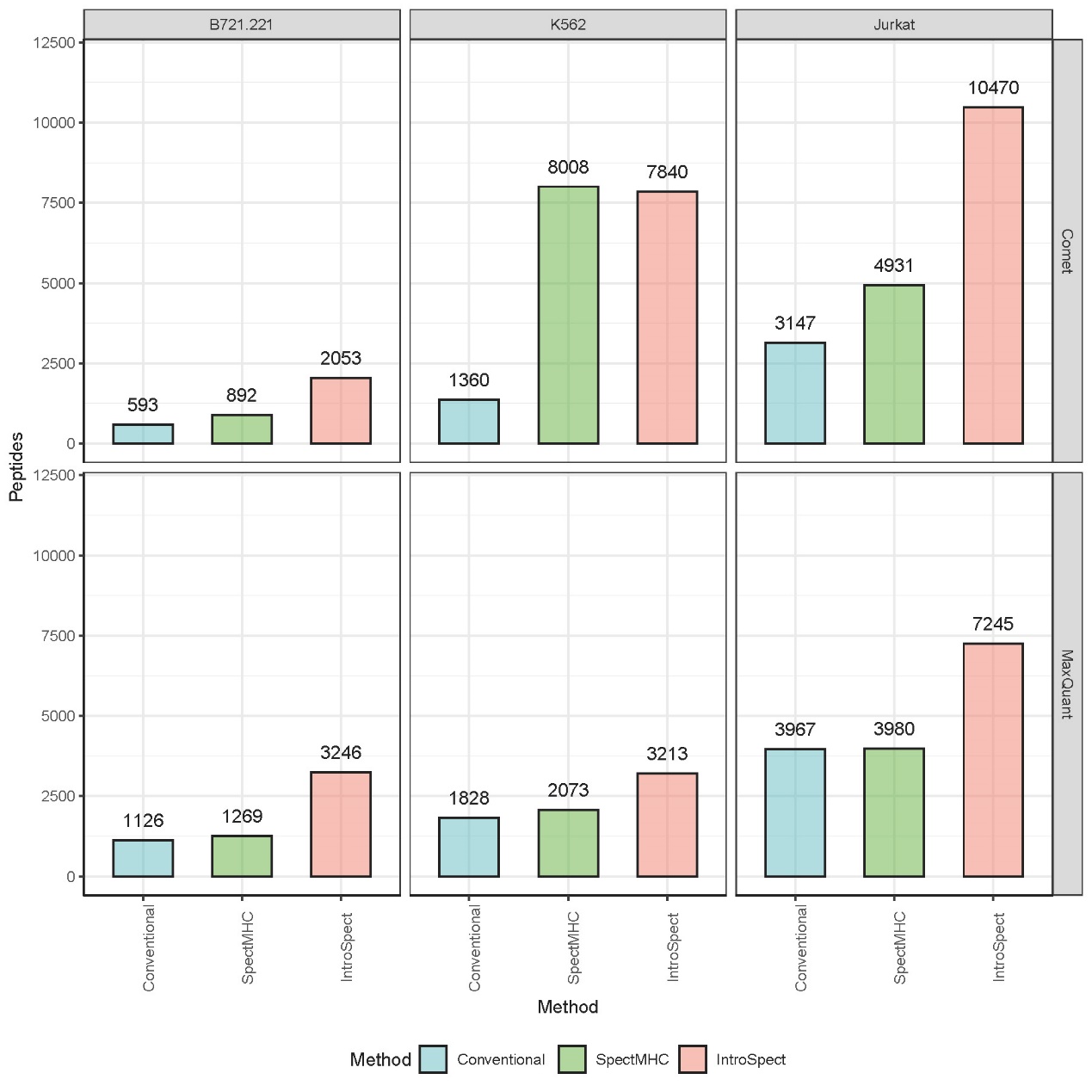

**Supplementary Figure 3.** The performance of IntroSpect on Comet and MaxQuant search tools.

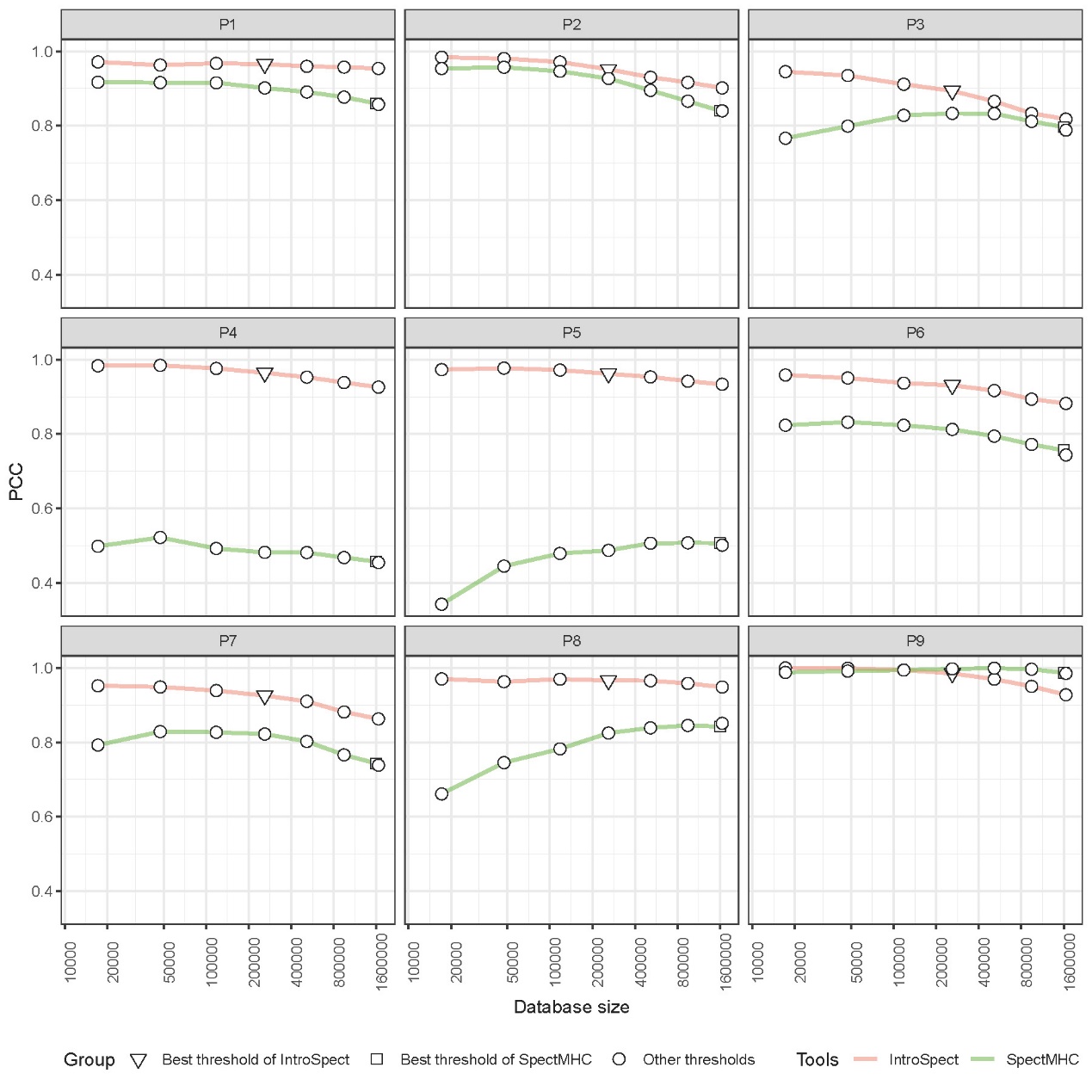

**Supplementary Figure 4.** The comparison of PCC_aaf_ on each position between the IntroSpect and SpectMHC.

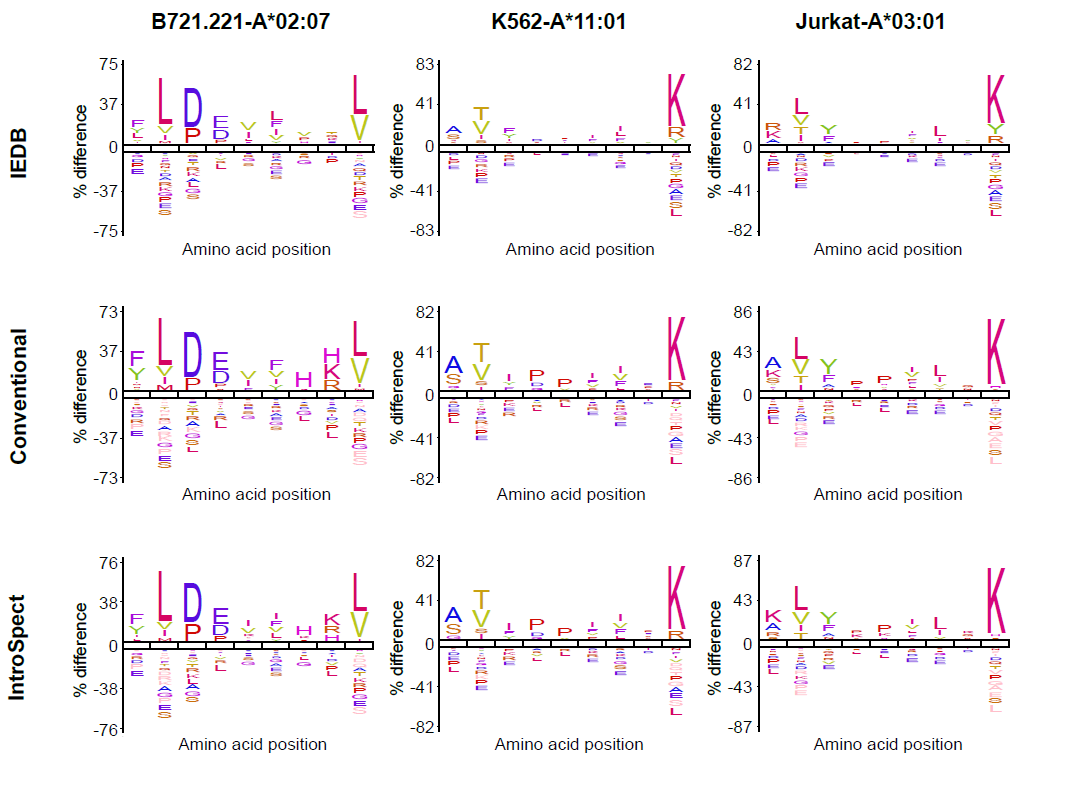

**Supplementary Figure 5.** The sequence logo comparison of immunopeptides in various datasets (B721.221-A*02:07, K562-A*11:01 and Jurkat-A*03:01) by the conventional search, IntroSpect search and IEDB downloaded.

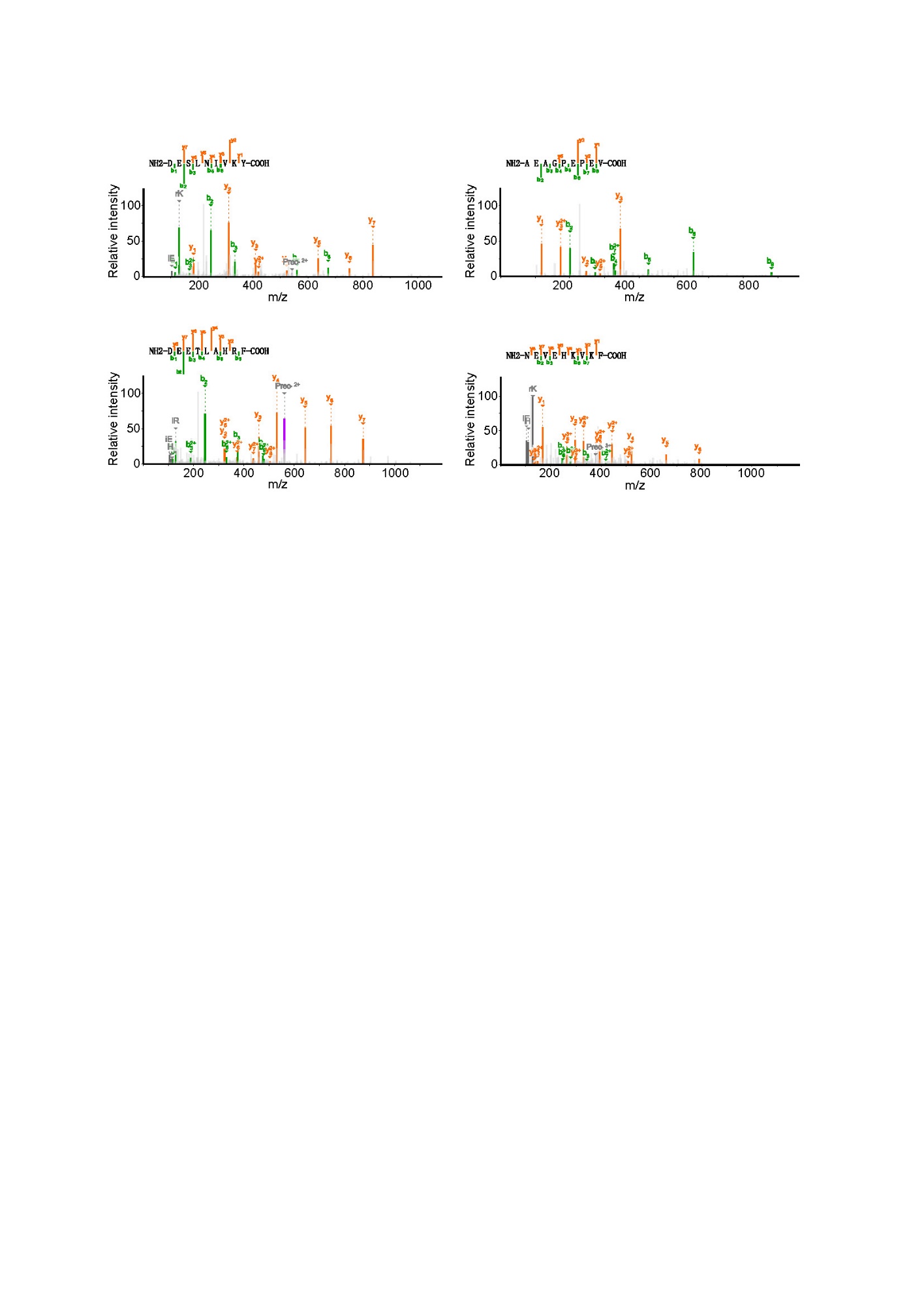

**Supplementary Figure 6.** Spectra of neoantigen candidates assigned by both the conventional search and IntroSpect search.

**Supplementary Table 1.** The neoantigens identified from HCT116 cell line.

| **ID** | **Peptide** | **Gene** | **Mutation type** | **Mutation site** | **Expression (TPM)** | **HLA allele** | **Predicted**  **IC50 (nM)** | **Validation** | **Identified from the Conventional (C) or IntroSpect (I)** |
| --- | --- | --- | --- | --- | --- | --- | --- | --- | --- |
| Neo-1 | AEAGPEPEV | EIF3B | SNV | p.S64P | 58.78 | B*45:01 | 279.57 | ligand presentation | Both |
| Neo-2 | VTEPGTAQY | AKAP13 | SNV | p.M452T | 11.61 | A*01:01 | 24.87 | IFNg release | I |
| Neo-3 | KLDAGEQRL | AMZ2 | SNV | p.N30D | 37.15 | C*05:01 | 93.65 | ligand presentation | I |
| Neo-4 | ASELHTSLY | MDN1 | SNV | p.H3423Y | 14.65 | A*01:01 | 9.43 | ligand presentation | I |
| Neo-5 | SLMEQIPHL | CKAP2 | INDEL | p.K603X | 18.67 | A*02:01 | 3.59 | qualitative binding;  ligand presentation;  IFNg release | I |
| Neo-6 | QTDQMVFNTY | CHMP7 | SNV | p.A324T | 12.87 | A*01:01 | 15.85 | ligand presentation | I |
| Neo-7 | DESLNIVKY | CCZ1B | SNV | p.E71D | 15.66 | B*18:01 | 15.83 |  | Both |
| Neo-8 | DEETLAHRF | CWF19L1 | SNV | p.R523H | 10.90 | B*18:01 | 39.72 |  | Both |
| Neo-9 | NEVEHKVKF | SYNE2 | SNV | p.I2942V | 11.92 | B*18:01 | 25.22 |  | Both |
| Neo-10 | AEADAALQKA | USH1C | SNV | p.E488D | 4.45 | B*45:01 | 32.88 |  | I |
